# Topological transitions, turbulent-like motion and long-time-tails driven by cell division in biological tissues

**DOI:** 10.1101/2022.11.25.518002

**Authors:** Xin Li, Sumit Sinha, T. R. Kirkpatrick, D. Thirumalai

**Affiliations:** Department of Chemistry, University of Texas, Austin, TX 78712, USA; Department of Physics, University of Texas, Austin, TX 78712, USA; lnstitute of Physical Science and Technology, University of Maryland, College Park, Maryland 20742, USA; Department of Chemistry and Department of Physics, University of Texas, Austin, TX 78712, USA

## Abstract

The complex spatiotemporal flow patterns in living tissues, driven by active forces, have many of the characteristics associated with inertial turbulence even though the Reynolds number is extremely low. Analyses of experimental data from two-dimensional epithelial monolayers in combination with agent-based simulations show that cell division and apoptosis lead to directed cell motion for hours, resulting in rapid topological transitions in neighboring cells. These transitions in turn generate both long ranged and long lived clockwise and anticlockwise vortices, which gives rise to turbulent-like flows. Both experiments and simulations show that at long wavelengths the wave vector (*k*) dependent energy spectrum *E*(*k*) ≈ *k*^−5/3^, coinciding with the Kolmogorov scaling in fully developed inertial turbulence. Using theoretical arguments and simulations, we show that long-lived vortices lead to long-time tails in the velocity auto-correlation function, *C*_*v*_(*t*) ∼ *t*^−1/2^, which has the same structure as in classical 2D fluids but with a different scaling exponent.

## I INTRODUCTION

The cell, an active agent, is the basic unit of life. The motion of a single cell has been studied intensively in the past few decades^1–8^. However, cells interact with each other and move in a collective manner, which plays an essential role in many biological processes, such as embryonic morphogenesis, organ development, wound healing, and spreading of cancer^9–11^. The collective movement of cells using physical models, spanning from sub-cellular to supra-cellular length scales^12–14^, gives rise to unexpected spatial and temporal correlations.

Since the discovery that flow-induced stretching of a long polymer in a high viscosity solution could produce flow patterns resembling that found in fully developed turbulence^15^, a number of studies have shown that turbulent-like features emerge in a variety of biological systems^16–20^. In these examples, the complex flow patterns are thought to rise from active forces, and hence the collective behavior is referred to as active turbulence, even though the scaling behavior is not universal. Hydrodynamic models, that build on the Toner-Tu equations^19–22^ and particle based simulations for self-propelled rods^20^ have generated turbulent-like patterns. In addition, numerical solutions and theoretical arguments of active nematic liquid crystal models have been used to show that vortex formation and turbulent-like behavior emerges^23–25^. In all these models, active forces, the cause for turbulent-like behavior, is introduced phenomenologically (by “hand” as it were).

Recent studies^18,26^ suggest that cell division and apoptosis lead to turbulent-like velocity fields with long-range vortex structures in two dimensional (2D) cell monolayers upon averaging over hundreds of division events. In addition, it is found that cell displacements exhibit super-diffusive behavior in a 2D confluent monolayer^27^, where cell division and apoptosis occur continuously. By building on our previous studies^28,29^, which showed that imbalance between the rates of cell division and apoptosis, leads to unusual dynamics in a growing tumor spheroid, we investigate if a similar picture could be used to analyze the time-dependent cell trajectories generated in the 2D monolayer^27^ to generate new physical properties.

Here, we investigate the collective cell migration induced by division and apoptosis in a confluent 2D cell monolayer using experimental data on Madin-Darby Canine Kidney (MDCK) cells^27^, in combination with an agent-based simulations^28,29^. We find that cell division leads to a directed motion of cells, which explains the super-diffusive behavior of cells with the mean-square displacement, Δ(*t*) ∝ *t*^1.5^. The directed motion of the new born cells induce rapid rearrangements of the neighbors in the confluent tissue through topological (T1 and T2 transitions) changes. The cell rearrangement induced by a single cell division leads to vortex formation. With the occurrence of multiple cell births and deaths, a turbulent-like behavior naturally emerges with the mean size of the vortices being ∼ 100 *μm*. Interestingly, we find two scaling regimes for the kinetic energy spectrum, *E*(*k*) with one following a Kolmogorov-like power law (*E*(*k*) ≈ *k*^−5/3^). The long-range correlated vortex motion leads to a long time tail (LTT) for the velocity auto-correlation function (*C*_*v*_(*t*)), which decays as *C*_*v*_(*t*) ∼ *t*^−1/2^, which differs from the well-known result 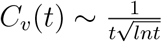 found in classical hard-disc-model fluids.

## II. RESULTS

### Vortex motion in a confluent cell monolayer

A confluent MDCK monolayer exhibits rich dynamics (see Figs. S1-S5 for some general features of the monolayer as it evolves in time) even though the cells are jammed. To probe the motility of the cells in the collective, we created the velocity vector field by applying the digital particle image velocimetry^30^ to the experimental data^27^. Collective motion of cells, in which a group of cells move in the same direction with similar magnitude of velocity (**v**)^31^, occurs in different regions of the monolayer (see Fig. 1a). Interestingly, formation of several vortices are discernible in the velocity field, which indicates a coherent motion of cells (see the dashed rectangle in Fig. 1a as an example). We calculated the vorticity field (***ω*** ≡ ∇ × **v**), which describes the local rotational motion of the velocity flow (see Fig. 1b). Positive vorticity (yellow color) indicates clockwise rotation, while negative vorticity (blue color) shows anticlockwise rotation. The vortices in Fig. 1a are clearly elucidated in the vorticity field (see the bright yellow circle in the lower left of Fig. 1b and the dashed rectangle in Fig. 1a). Outward and inward velocity fluxes, characterized by divergence (div(**v**) ≡ ∇ · **v**) of the velocity flow (Fig. 1c), are non-uniform in the monolayer.

**FIG. 1:**
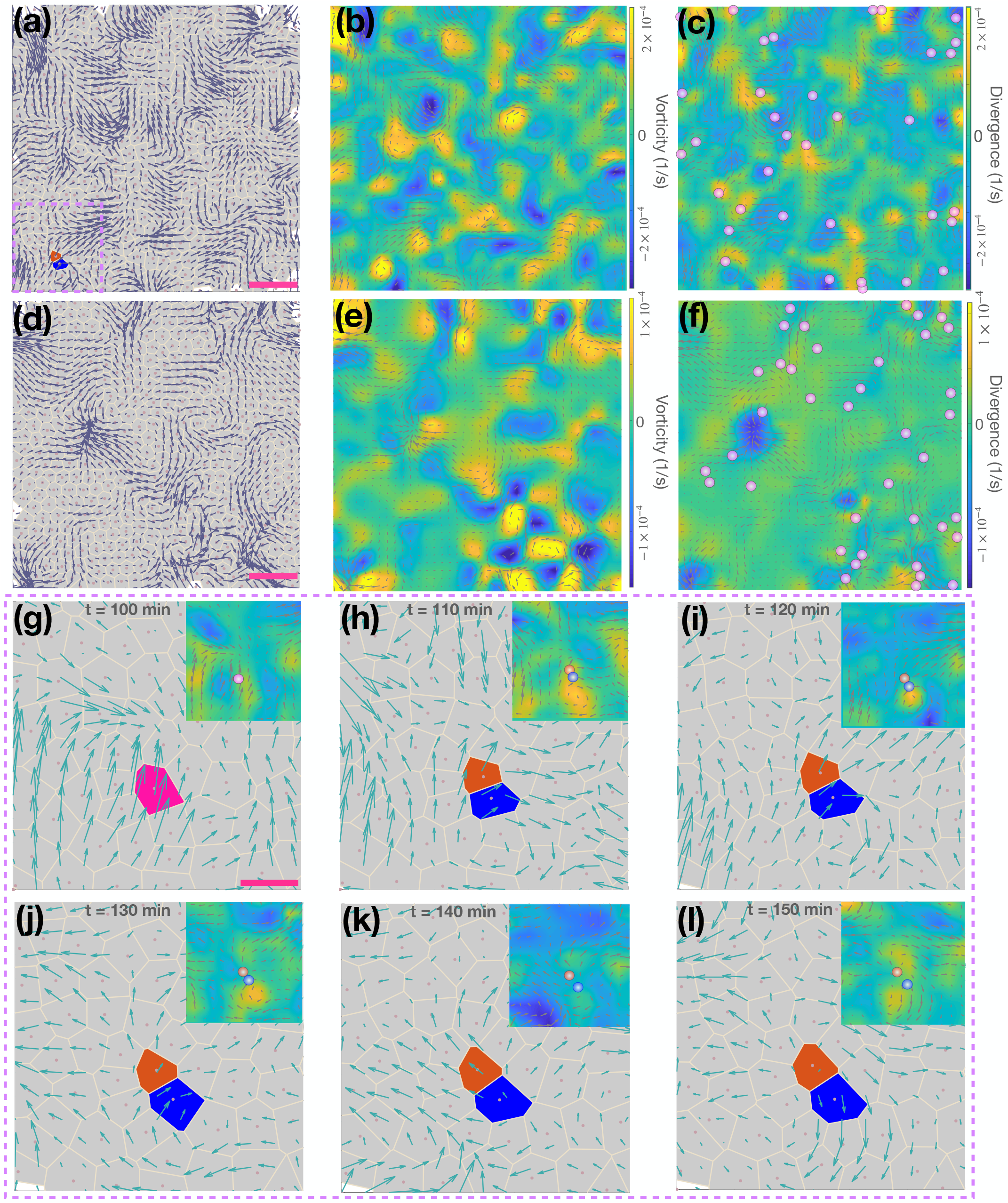
Experiments and simulations (an agent-based model) on cell motility in a MDCK monolayer. **(a)** A typical velocity vector field in a MDCK cell monolayer, at time *t* = 120 min, illustrated by the blue arrows. The positions of the cell nuclei are indicated by the pink dots from which the Voronoi cells are constructed. **(b)** The vorticity of the cell monolayer for the same time frame as in Fig. 1(a). The magnitude and direction of the rotation of the vorticity are coded by the color bar on the right side of the figure. **(c)** The divergence field of the cell monolayer at *t* = 120 minute. The magnitude of the divergence is color coded, with outward flux in yellow and inward flux in blue (see the color bar on the right side of the figure). The purple dots show the positions of the new cells, which are born between time *t* = 95 minute to *t* = 120 minute. The velocity vector field is overlaid on the vorticity and divergence fields. Same as **(a-c)**, the velocity vector **(d)**, vorticity **(e)**, and divergence **(f)** fields calculated from simulations. The pink bar in **(a)** and **(d)** corresponds to 50 *μm*. The same time interval (25 minutes) as in **(c)** is used in **(f)** for the positions of the new cells. **(g-l)** A zoom in of the dashed rectangle region at the lower left corner of Fig. 1**(a)** for different times, which shows the formation of a vortex accompanied by a single cell division event. Cell division is observed in Fig. 1**(h)** (see the orange and blue cells originated from the pink cell division in Fig. 1**(g)**), and a vortex is formed that accompanied by such an event (see the velocity field close to the blue cell). To illustrate the formation of the vortex vividly, the vorticity field, as in Fig. 1**(b)**, is also shown in the inset of each figure at the upper right corner (see the yellow regions in Fig. 1**(h-j)**, especially). The positions of the mother and daughter cells are indicated by dots in different colors. The pink bar in **(g)** corresponds to 20 *μm*.

### Cell division as the driver of vortex motion

Previous theoretical and experimental studies^28,32–34^ have shown that active forces generated by cell division drives fluidization of tissues, leading to heterogeneous cell motility patterns both in a two-dimensional monolayer and in the three-dimensional spheroids. Such events (Fig. S1c) occur frequently in the experiments, as shown in Figs. 1a-c. Therefore, we surmised that the complex flow patterns observed here are driven by cell birth and death. To test this notion, we first show the positions of newly generated cells (born about 30 minutes before the snapshot in Fig. 1a) in the divergence field (see the purple dots in Fig. 1c). The origin of the outward flux (colored in yellow) is frequently accompanied by the appearance of newly created cells nearby. In addition, new cells arising from cell divisions are close to the vortices (see Figs. 1b-c). Therefore, it is likely that the flow patterns (Figs. 1a-c) in the monolayer are driven by active forces produced by the cell division.

### Particle based simulations

To further elucidate the relevance of cell division, we simulated the dynamics of a cell monolayer using an agent-based model (details are in the Methods section and the SI). The velocity flow pattern in the simulations and experiments are similar (compare Figs. 1a and 1d). In addition, several pairs of vortices with opposite direction of rotation are also observed (see the lower middle of Fig. 1e). Note that in contrast to other studies^35–37^, we do not include self-propulsion in our model. Thus, the observed velocity flow and vortex formation can only arise from self-generated active forces (SGAFs) arising from cell division. Indeed, cell division (see the purple dots in Fig. 1f), which produces an outward flow in the the divergence field (Fig. 1f), is close to the vortex location (see Figs. 1d-e). Hence, cell division and apoptosis generate complex cell motility patterns, resembling turbulent-like structures (Figs. 1a-f).

To illustrate how cell division leads to vortex formation, we tracked the collective cell motion in the dashed rectangle region in Fig. 1a for one hour (see Fig. 1g-l). The mother cell (in pink, see Fig. 1g at *t* = 100 min) divides into two daughter cells (orange and blue) at *t* = 110 min (see Fig. 1h). The vortex observed in Fig. 1a-b at *t* = 120 min is established at *t* = 110 min (see both the velocity and vorticity fields in Fig. 1h) immediately after cell division. The vortex persists for an additional 20 minutes (Fig. 1i-j) before vanishing (Fig. 1k-l). In contrast to previous experimental observations^18,26^, where vortices are discernible only upon averaging over 100 division events, we found that a single cell division event leads to a vortex, lasting for more than 30 minutes. In addition, multiple cell division events drive a turbulent-like flow (Fig. 1a-f).

### Directed motion of new born cells leads to anomalous diffusion

To study how vortices are formed by cell division explicitly, we started by monitoring the cell dynamics in the trajectories during the first 100 minutes for the newly created cells (see Fig. 2a). For illustration purposes, we set the initial positions of all the cells at the origin. There is no obvious preferred direction of motion for cells, which suggests that they move isotropically (Fig. 2a). Surprisingly, the motion is directed after cell division over a distance of ∼ 10 *μm* (see Fig. 2a and the inset figure for a few representative trajectories). The simulations also show a similar directed motion of cells during the first 100 minutes after cell divisions (see Fig. 2b).

**FIG. 2:**
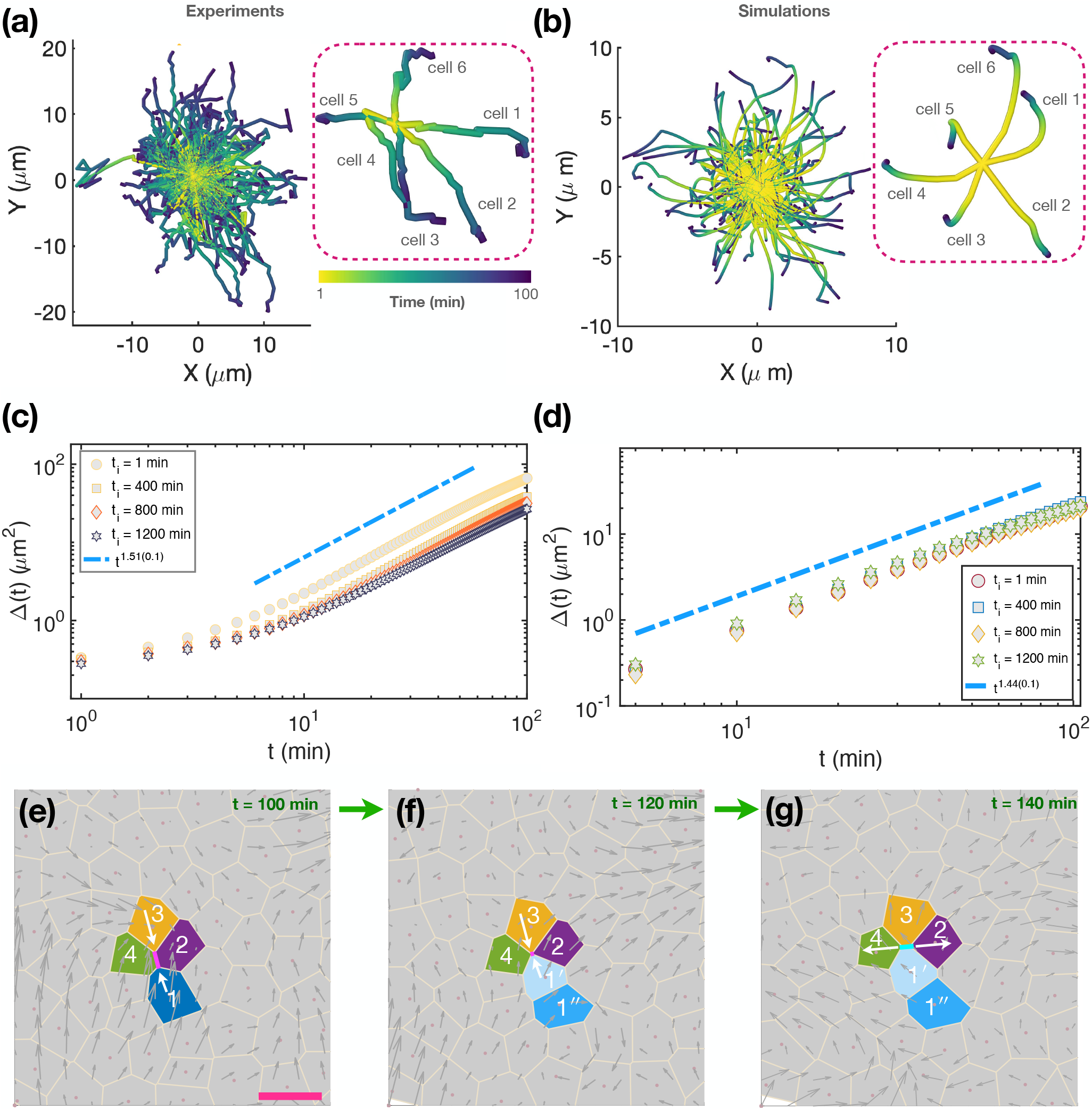
Anomalous diffusion and topological changes. **(a)** The trajectories of the first 150 newly generated cells for the initial 100 minutes after birth. The initial position of the new cells are aligned at the origin (0,0) for illustration purpose only. Inset on the right: Six randomly chosen representative trajectories from **(a). (b)** Same as **(a)**, except they are from simulations. The trajectories in **(a)-(b)** are color coded by time (scale on the bottom of the inset in **(a)**). **(c)** Mean-square displacement (Δ(*t*)) of cells calculated over a time window of 200 minutes, starting from different initial time points *t*_*i*_ over the course of the experiment^27^. **(d)** Same as **(c)**, except the results are from simulations. The same time window of 200 minutes is used for each *t*_*i*_. The slope (1.44) of the dashed-dotted line from simulations is similar as shown in **(c)** for experiments (1.51), both showing superdiffusive behavior. **(e)-(g)** T1 transition induced by cell divisions. Cell 1 in **(e)** divided into two cells (cell 1′, and cell 1′′) in **(f)**. The motion of newly generated cells leads to T1 transition, which separates the two neighboring cells (cell 2 and cell 4) away from each other in **(g)** (see also Fig. S7 in the SI for more details). The pink bar in **(e)** corresponds to 20 *μm*. Same as Figs. 1g-l, a vortex around cells 1′ and 1′′ is found in **(f)** (see the velocity vector field in grey arrows).

To illustrate the consequence of cell-division induced directed motion, we calculated the mean-square displacement (Δ(*t, t*_*i*_) = ⟨[**r**(*t*_*i*_ + *t*) − **r**(*t*_*i*_)]^2^⟩) of cells at different initial times *t*_*i*_, where **r** is the cell position. Both experiments and simulations (Figs. 2c-d) exhibit superdiffusive behavior, Δ(*t, t*_*i*_) ∝ *t*^*α*^, with an exponent *α* ≈ 1.51 ± 0.1 and 1.44 ± 0.1 respectively, irrespective of the time windows used. The displacement distributions, (*P* (Δ*x*), *P* (Δ*y*)), of cells calculated from both experiments and simulations (see Fig. S6a-d in the SI) deviate from the Gaussian distribution (see the green solid lines). It follows that cell division and apoptosis is the origin of superdiffusive motion of cells.

### Cell division and apoptosis result in topological changes

The SGAFs^38^ due to cell division and apoptosis lead to a directed motion of newly generated cells, which influences the movement of cells nearby in a confluent tissue. Interestingly, there are several T1 transitions, which are critical during morphogenesis^39^, and cancer invasion^40^, during a single cell division in the MDCK cell experiments. One example is shown in Figs. 2e-g (see also Fig. S7 in the SI). The force produced by the division of cell 1 at *t* = 110 min separates the two neighboring cells (cells 2 and 4) away from each other but brings cells 1 and 3 together at later times. Such rearrangement of cells 1 − 4 and other cells nearby finally leads to vortex formation (see the velocity vector field in Figs. 2e-g and also Figs. 1g-l).

In addition to a single vortex induced by one cell division, (Figs. 1g-l), there is a pair of vortices, with opposite sense of rotation, in certain time frames (see the dashed rectangle in Fig. 3a). In the same region, there are three cell divisions, which lead to dramatic cell rearrangements nearby, 10 minutes before a vortex pair formation (the purple dots in the the dashed rectangle). To describe the formation of one vortex pair quantitatively, we plot the vorticity value along the arrow in the dashed rectangle in Fig. 3a using the digital particle image velocimetry^30^. A smooth transition from anticlockwise to clockwise vorticity is clearly depicted (see Fig. 3b). Similarly, several pairs of vortices with opposite directions of rotation are found in simulations (see the lower middle of Fig. 1e). Besides, there is evidence for T2 transition (Fig. S8 in the SI), which arises from the extrusion (apoptosis) of a cell from the confluent monolayer. Taken together, these results suggest that cell division and apoptosis not only regulate cell numbers during morphogenesis and cancer development, they also drive dramatic cell rearrangements, as discovered recently for chick gastrulation^41^, and germband extension of Drosophila^42^. Finally, the cell rearrangement through topological T1/T2 transitions leads to the formation of vortices and turbulent-like structure in the confluent MDCK tissue.

**FIG. 3:**
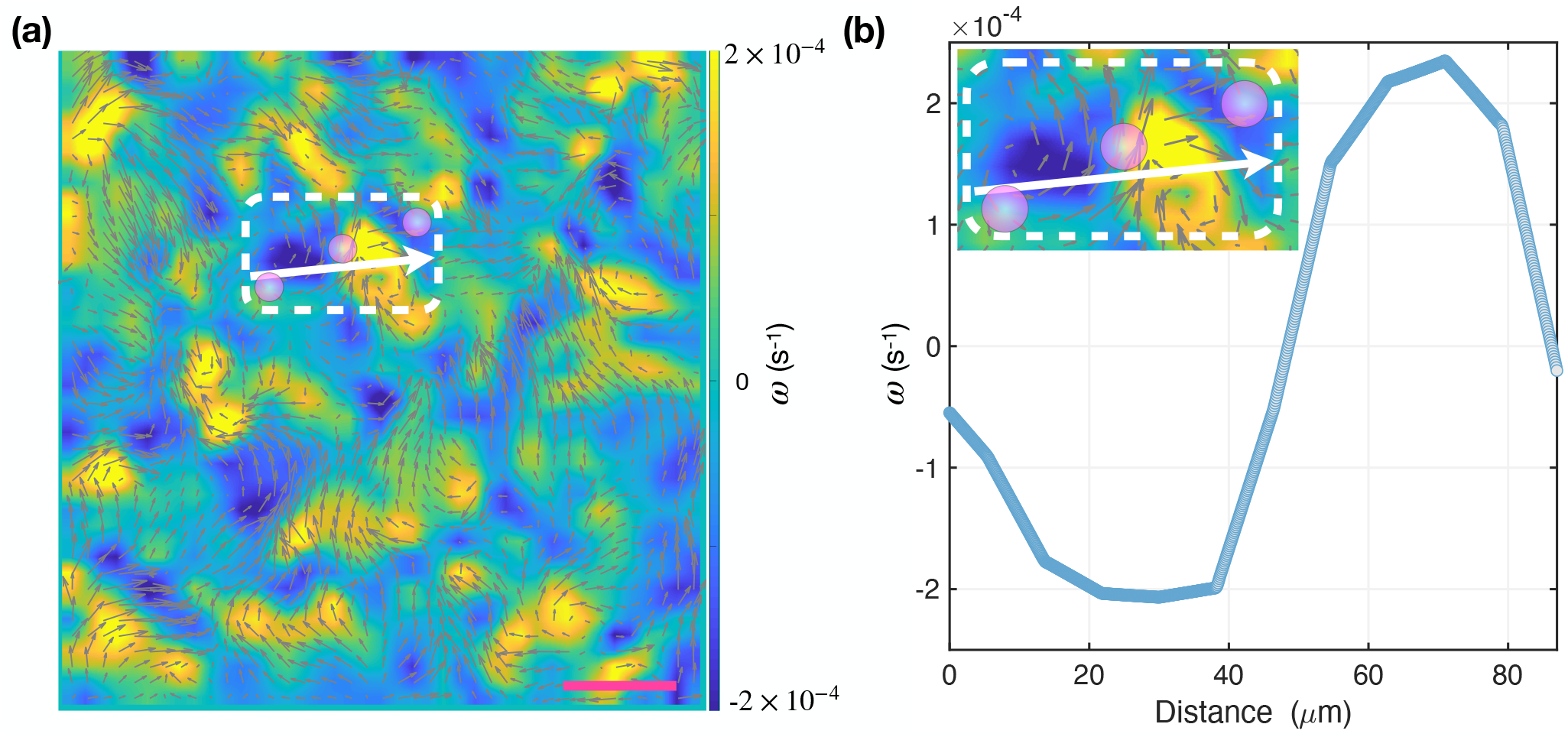
Clockwise and anticlockwise vorticity (*ω*). **(a)** A pair of vortices with clockwise and anticlockwise rotations at time *t* = 80 minute using the vorticity field of a cell monolayer (see the region indicated by the dashed rectangle). Three purple circles mark three cells, which were born between *t* = 70 − 80 minutes. The velocity vector field is overlaid on the vorticity fields. The pink bar corresponds to 50 *μm*. Scale on the right gives *ω* in unit of *s*^−1^. **(b)** Vorticity as a function of the distance from the left side of the dashed box along the white arrow in **(a)**. A zoom in of the dashed rectangle in **(a)** is shown in the inset.

### Spatial correlations

To probe the spatial variations in the turbulent-like flow, we calculated space-dependent correlations associated using the velocity field, *C*_*v*_(*R*). From the cell position, **r**_*i*_(*t*), of the *i*^*th*^ cell at different times, we calculated the velocity, **v**_*i*_(*t*) ≡ [**r**_*i*_(*t* + *δt/*2) − **r**_*i*_(*t* − *δt/*2)]*/δt*, at time *t* where *δt* is the time interval. The spatial velocity correlation function (*C*_*v*_(*R*)) is defined as,

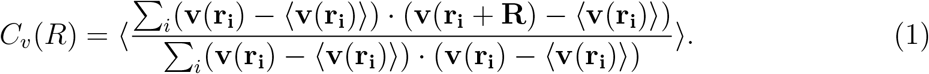

The correlation function at different times (see Fig. 4a) has negative minimum at a distance *R*_*v*_ ≈ 100 *μm*, which is about 10 times greater than the mean cell size (see Fig. S3). The negative values for the correlation function are due to the anti-parallel velocity on the opposite sides of the vortices (see Fig. 1b). Therefore, *R*_*v*_ is roughly the size of the vortices (Fig. 1), which are about 100 *μm*. The *C*_*v*_(*R*) calculated from our simulations supports such a link (see Fig. 4b). Interestingly, the collective motion of cells is quite robust, persisting for a long time, despite the increase in the cell density (Fig. S4a).

**FIG. 4:**
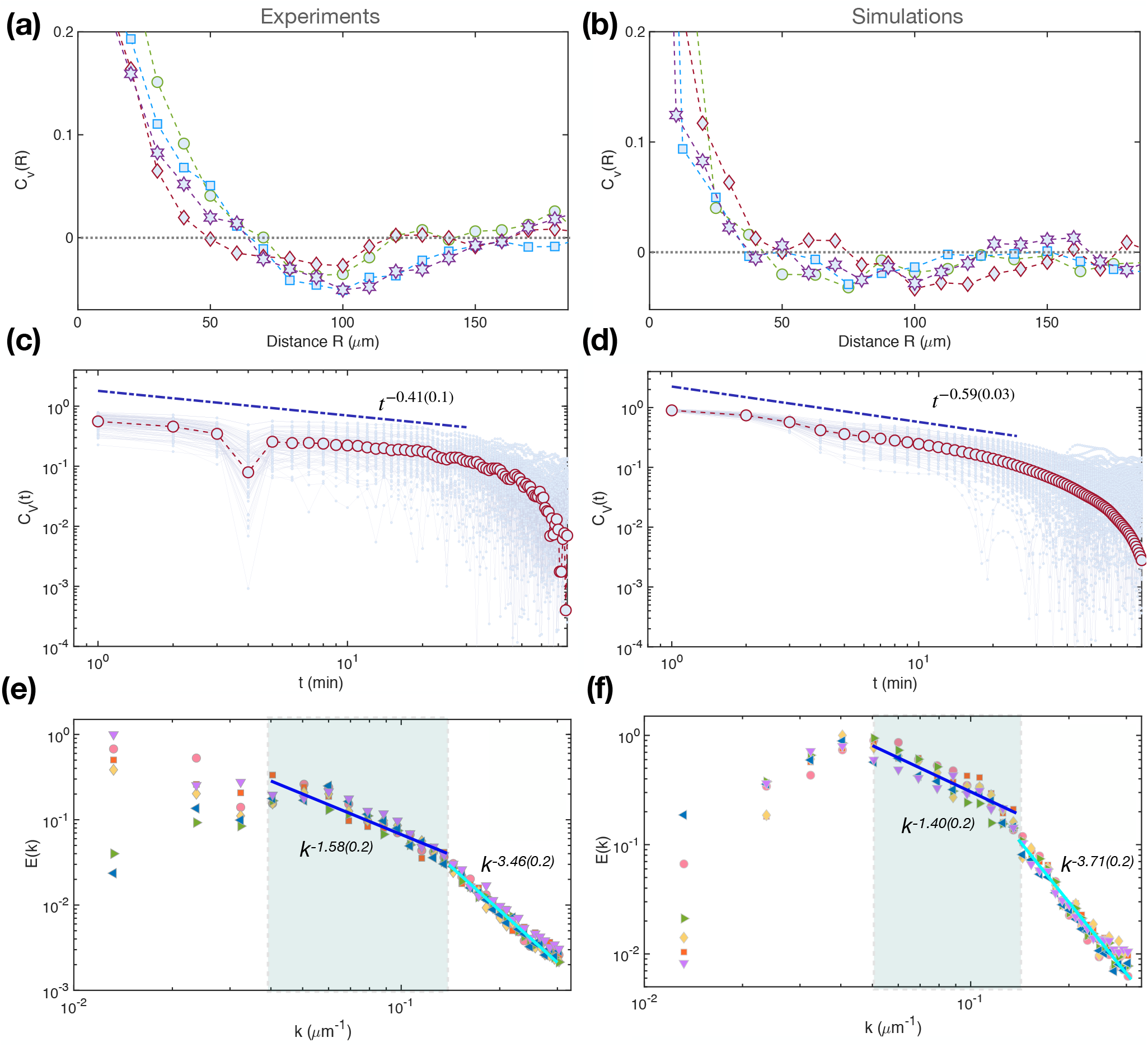
Velocity autocorrelation and energy spectra. **(a)** The velocity spatial correlation (*C*_*v*_(*R*)) at different times: *t*_*i*_ = 100 (cirlces), 600 (squares), 1200 (diamonds), and 1400 (hexagrams) minutes. Each curve is averaged over 10 time frames. A negative minimum value at *R*_*v*_ ≈ 100 *μm* (≈ 10 times the cell size) is present in all the curves. **(b)** Same as **(a)**, except it is obtained from simulations. We took *δt* = 10 minutes in **(a-b)** and the dotted lines show the zero value. **(c)** The velocity autocorrelation (*C*_*v*_(*t*)) as a function of time in a double logarithmic scale calculated using *δt* = 4 minutes (see Fig. S12 in the SI for different values of *δt*). The data from a single cell analysis (135 cells in total which are alive through the whole experiment) is shown in grey dots and the red circles show the mean value. The slope of the dash-dotted line is -0.41±0.1. **(d)** Same as **(c)**, except the result is obtained from simulations. The slope of the dashed-dotted line is -0.59±0.03. **(e)** The energy spectra *E*(*k*) calculated using the trajectories generated in imaging experiments at different times: *t* = 100 (cirlces), 110 (squares), 120 (diamonds), 130 (right-pointing triangle), 140 (left-pointing triangle) and 150 (down-pointing triangle) minutes. **(f)** Calculated *E*(*k*) from simulations. The values of the slope of the solid lines are shown in the figure. In the grey regime *E*(*k*) ∼ *k*^−1.58^ from experiments. This value (1.58) is hauntingly close to 5/3 expected for inertial turbulence. The two exponent values obtained from simulations (1.4 and 3.7) are close to the values obtained from analysis of the data from experiments (1.6 and 3.5). The unit for the wave vector (*k*) is *μ*m^−1^, and *E*(*k*) (*μ*m^3^/s^2^) is scaled by the maximum value.

### Long time tails (LTT)

We calculated the velocity correlation function 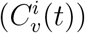 for cell

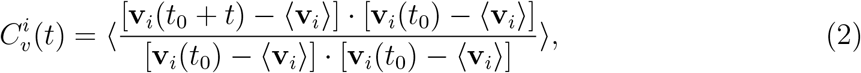

where ⟨**v**_*i*_⟩ is the mean velocity of cell *i*, and the average is over time *t*_0_ (24 hours). The ensemble average *C*_*v*_(*t*) for the monolayer is obtained by averaging 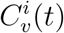 over all the cells. The decay of *C*_*v*_(*t*) follows *C*_*v*_(*t*) ∼ *t*^−*β*^ with *β* = 0.4 ± 0.1 (see the blue dash-dotted line in Fig. 4c). Our simulations also show a power-law decay with *β* = 0.59 ± 0.03 (see Fig. 4d). Based on dimensional argument, we expect that the decay of *C*_*v*_(*t*) ∼ Δ(*t*)*/t*^2^ ∼ *t*^−0.5^, which is fairly close to the experimental and simulation results. Using theory that accounts for SGAFs, the exponent *β* = 1/2 in 2D (see the Methods section). A comparison from three different fit functions to the experimental and simulation data are also shown in the Fig. S9 in the SI. The emergence of LTT in *C*_*v*_(*t*) shows that motion of cells is persistent in a specific direction (Figs. 2a-d), which in turn is linked to vortex motion. To ascertain whether the power-law relation depends on the time interval selected to calculate *C*_*v*_(*t*), we varied *δt*. The value of *β* stays around 1/2 (see Fig. S10 in the SI).

A long time tail relation is also found for the current-current correlation functions in disordered systems, which decays by *t*^−*d/*2^ (*d* is the dimension of space)^43^. Such a relation was first reported in simulations of classical hard-disc (sphere)-model fluids^44^, and was explained by correlated collision events theoretically^45,46^. The difference in the value of *β* between abiotic systems and MDCK monolayer is due to the generation of SGAFs due to cell division, which results in persistent (almost ballistic) directional motion of cells for long times.

### Energy spectrum

It is clear that turbulent-like flow emerges naturally in the collective motion of cells (Figs. 1a-c). To characterize the nature of turbulent-like motion in the epithelial cells, we calculated the wave vector (*k*) dependent energy spectrum, *E*(*k*), at different times (see Fig. 4e). There are two scaling regimes in *E*(*k*) as a function of *k*. In the intermediate values of *k, E*(*k*) ∼ *k*^−1.58(0.2)^. The value of the exponent is close to the Kolmogorov-Kraichnan prediction, −5/3, found in the inertial turbulence^47^. We found a similar exponent value (−1.4 ± 0.2) from our simulations (see Fig. 4f). In the MDCK cells, the Reynolds number is small (≈ 0)^48^, which shows that the underlying mechanisms must be quite different for the turbulent-like motion^16^.

At smaller scales (large *k*), the results (Fig. 4e) obtained by analyzing the experimental trajectories show that *E*(*k*) ∼ *k*^−*λ*^ (*λ* = 3.5 ± 0.2). From the agent-based simulations, we deduce that *E*(*k*) ∼ *k*^−3.7(0.2)^. Two comments are worth making: (*i*) The *λ* values from experiments and simulations are fairly close. It is interesting that scaling anticipated in the context of inertial turbulence is obeyed in living systems, given that energy injection, which is a consequence of cell division and apoptosis, is autonomous. (*ii*) The prediction for inertial 2D turbulence at large *k* is *λ* ≈ 3, which differs from the estimated value for MDCK cells and simulations. It is unclear if the large *k* finding in living tissues is simply a consequence of non-universal behavior or if large *k* limit is not accessed in experiments and simulations. In active systems, such as microtubule-kinesin complex and bacterial suspensions, *λ* ranges from 2 to 4.5^16,20,49,50^.

## III. DISCUSSION

We investigated the dynamics of cell rearrangement and collective motion in a biologically relevant tissue^9–11^, by analyzing data from confluent MDCK cell monolayers, combined with agent-based simulations and theory. Cell division and apoptosis lead to a directed motion of cells for hours, which results in a universal super-diffusive behavior of cells characterized by the mean-square displacement exponent, *α* ≈ 1.51, irrespective of the initial time *t*_*i*_ considered in the experiments^27^. Our agent-based simulations yield *α* = 1.44 ± 0.1. It is worth emphasizing that, in the absence of cell division, there is complete cessation of motion in the simulations. Therefore, the complex dynamics in the simulations, which capture the salient features in the MDCK cells, arises solely due to self-generated active forces generated by cell division and apoptosis^28,29,38^.

The directed motion of newly generated cells induces rapid rearrangements of cells nearby through T1–cell intercalation and T2 transitions. Interestingly, cell rearrangements due to a single cell division result in vortex formation in confluent tissues, with lifetimes lasting for 30 minutes. More importantly, turbulent-like flux patterns emerges in the region where multiple divisions occur, which is also supported by our agent-based simulation model. From the spatial velocity correlation function, we estimate that the mean size of the vortices can be as large as to 100 *μm*, which is nearly 10 times larger than the size of a single cell (∼ 10 *μm*). Finally, the kinetic energy spectrum (*E*(*k*)) exhibits two distinct scaling behaviors of the collective cell movement. At intermediate values of *k, E*(*k*) ∼ *k*^1.58*±*0.2^, which is close to the predicted Kolmogorov-Kraichnan behavior for the inertial turbulence. At large *k* value, *E*(*k*) ∼ *k*^−3.5^, which does not seems to have counter part in inertial turbulence, and is likely to be non-universal. The exponent characterizing large *k* behavior of *E*(*k*) show substantial variations, depending on the system^16,51,52^.

In classical hard-disc-model fluids^44–46^, a long time tail is expected for the velocity autocorrelation function with 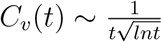. Both experiments and simulations show a slower power law decay *C*_*v*_(*t*) ∝ *t*^−1/2^ in the MDCK cell monolayer. To explain this finding, we developed a theory that accounts for active forces leading to the prediction of a slower decay, *C*_*v*_(*t*) ∝ *t*^−1/2^. The theory and the agent-based simulations indicate that self-generated active forces lead to highly correlated responses. Given the simple model considered here, we expect that our studies can be used to describe collective motion of other cell types or a mixture of different cell types, and even other active systems in general.

In addition, we showed that the amplitude for the mean-square displacement (Fig. 2c) and the mean velocity of cells (Fig. S4b) decrease, as the initial time *t*_*i*_ increases, which indicates a slowing down due to jamming or aging of the cell dynamics. Similar aging behavior has been noted in other types of cell monolayers^53^. In addition, all the dynamical characteristics for cells, (superdiffusion, compressed exponential relaxation and aging) are found in many soft glassy systems^54–56^ in the absence of active forces. Thus the physics used to describe abiotic glassy materials could be applied to understand living active matter^57,58^. Finally, more detailed models by incorporating the size reductive divisions^59^, maturation and strengthening of cell-cell adhesion^53^ or other potential processes could be developed based on our present study to explain aging and other complex cell activities.

## IV. METHODS

### Summary of MDCK epithelial cell experiments^27^

The MDCK (Madin-Darby Canine Kidney) epithelial cells were maintained in Dulbeccos Modified Eagle Medium, supplemented with 5% fetal bovine serum and 1% L-Glutamine. First, they were seeded in a 6-well plate to grow in complete medium until a confluent cell monolayer formed. Then, the images of the cell monolayer were acquired by Olympus IX81 inverted microscope (both in phase contrast and wide-field fluorescence) every minute for a 24-hour period. The phase-contrast microscopy was used to visualize the cell membrane and different organelles (see Fig. S1a). The nuclei of the cells at each time frame were also captured by wide-field fluorescence microscopy of EGFP-H2B expressing cells (see the cell nuclei shown in Fig. S1b for an example) and their positions were recorded from the single particle tracking by using the ImageJ^60^. A number of cell division events were observed in such a confluent monolayer (see Fig. S1c), while cell apoptosis/extrusion process led to the removal of cells from the monolayer. From the cell nuclei positions at each time frame, the cell trajectories can be obtained (see Fig. S1d, shows the first 150 minutes of recording for one of the field-of-views (FOV)). The cell trajectory in the figure exhibits a highly heterogeneous behavior. Some cells move in a rather straight and regular trajectory, while others exhibit irregular curly motion. The diverse movement is even clearer if the cell trajectories are plotted over the whole 24-hour period (see the two examples shown in the insets of Fig. S1d). The dynam-ically heterogeneous behavior is reminiscent of supercooled liquids, as was first shown by Angelini et al.^61^ in the context of cell monolayers.

### Particle-based simulations

We used a two-dimensional version of a particle-based model^28,29^ to simulate cell dynamics. We mainly focused on investigating the roles of the cell division and apoptosis on the collective cell motion in order to provide microscopic insights into experiments^27^. A single cell is modeled as a two-dimensional deformable disk. We use a periodic boundary conditions to mimic large systems. The cells interact with each other through repulsive (Hertzian contact mechanics) and attractive interactions (cell-cell adhesion). The motion of cells is described by an overdamped Langevin equation. The cells were allowed to grow and divide, as in the experiment. We also removed the cells from the monolayer at a constant rate to model the cell apoptosis and extrusion events that were observed in experiments. Because we do not consider self-propulsion, the cell motility can only arise from the self-generated active forces through cell growth, division and apoptosis^38^. The SI contains details of the model.

### Theory of LTT driven by cell division

The turbulent-like motion in the MDCK cells in which active forces are generated by cell division, suggests that we use methods developed to treat turbulence in conventional fluids^62–65^. The fundamental difference is that the random stirring that causes the turbulence in the conventional case is replaced by the internally generated cell division process in the biological case. Our aim is to compute the turbulent renormalized transport coefficients in the biological case, and from that determine the diffusive velocity autocorrelation function and the mean-squared-displacement. The relevant spatial dimension is *d* = 2.

The model dynamical equations are,

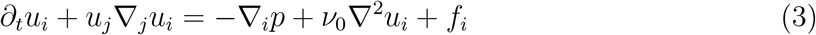

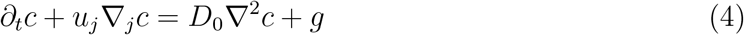

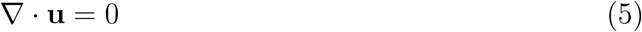

Here, *u*_*i*_ is the fluid velocity in the *i*-direction (repeated indices are to be summed over), *p* is the pressure, *ν*_0_ is the bare kinematic viscosity, *c* is the density of the active matter (MDCK cells), *D*_0_ is a bare diffusion coefficient. The equations given above couple vorticity in the fluid motion to cell density. We have also imposed the incompressibility condition in Eq. (5).

The statistical forces *f*_*i*_ and *g* are Gaussian noise terms that have zero mean and a nonzero two-point correlation functions that are local in space and time with gradient independent correlation strengths given by Δ_*f*_ and Δ_*g*_, respectively.

We treat the nonlinearities in these equations as a perturbation and work in Fourier space in both position and time, (**k**, *ω*). Further, the equations for both *u*_*i*_ and *c* are diffusive ^1^ and because *f*_*i*_ and *g* have identical statistical properties, we focus only on the mathematical structures and scaling solutions. We refer to the bare transport coefficients *D*_0_, and their renormalized values by *D*. Structurally the one-loop contribution to *D*, denoted by *δD*, is given by,

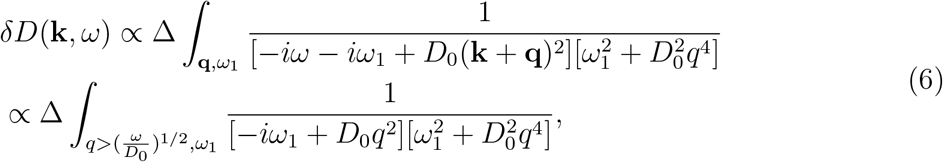

where we have assumed that Δ ∝ Δ_*f*_ ∝ Δ_*g*_. In going from the first to the second line in Eq. (6) we have replaced the external frequency and wavenumber dependence in the integrand by a frequency cutoff. For **k** = 0, this can be justified in a scaling sense.

Carrying out the frequency and wave number integrals in *d* = 2 gives,

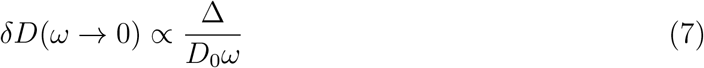

Since *δD* diverges as *ω* → 0, we use *δD* ≈ *D* and self-consistently replace *D*_0_ in Eq. (7) by *D*. We conclude that in *d* = 2,

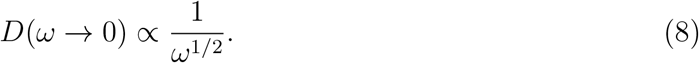

This result in turn implies that the velocity autocorrelation function behaves as *C*(*t* →∞) ∝ 1*/t*^1/2^ and the mean-squared displacement is ∝ *t*^3/2^. These results are consistent with our numerical work as well as with experiments. In a similar manner, one obtains 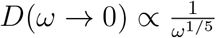, which implies a velocity auto-correlation function ∝ 1*/t*^4/5^ in *d* = 3.

## Acknowledgements

We are grateful to Giorgio Scita et al. for providing us the original experimental data. We thank Davin Jeong for help in producing figures. This work is supported by the National Science Foundation (PHY 17-08128), and the Collie-Welch Chair through the Welch Foundation (F-0019).

## Competing financial interests

The authors declare no competing financial interests.

## Footnotes

1 The pressure term in Eq. (3) is only used to enforce the incompressibility condition given by Eq. (5).

## Notes

### Competing Interest Statement

The authors have declared no competing interest.

