## Supplementary Information for "Topological transitions, turbulent-like motion and long-time-tails driven by cell division in biological tissues"

(Dated: 25 November 2022)

[a)] Electronic mail:

In the Supplementary Information (SI), we provide a summary of experimental results, several additional findings, description of the simulations using a particle-based model, and related discussions that are pertinent to the results present in the main text.

**Microscopy Experiments:** Because the present study uses results from the imaging experiments[1], we describe them in some detail. Two types of microscopes, phase-contrast and wide-field fluorescence microscopy are used in the MDCK cell experiments. A confluent monolayer, with cells distributed evenly over the whole space, is imaged using phase contrast microscopy, allowing the capture of the cell membrane (see Fig. S1a and movie 2 in the supplementary materials of the original article[1]). In addition, Enhanced Green Fluorescent Protein (EGFP)-tagged H2B histones were constitutively expressed by retroviral infection using standard protocols[1]. The images of the nuclei of all the cells, at each time frame, were captured by the wide-field fluorescence microscopy (see Fig. S1b and also the movie 1 in the supplementary materials in the experiments[1]). Cell division occurred frequently (see Fig. S1c). The left panel of Fig. S1c shows an example of a cell division event using the phase-contrast microscopy, while the right panel illustrates the same event using the wide-field fluorescence microscopy. The protocol in the experiments[1] was designed to avoid any directional bias for cell motility patterns in order to reduce the inhomogeneity and anisotropy of the cell density and shape in the monolayer. Using MosaicSuite plugin of the ImageJ[2,3] for single particle tracking over the fluorescence images of the cell nuclei (one-minute resolution over 24 hours), the positions of all the cells were recorded in consecutive frames. Finally, the trajectories of all the cells in one field-of-view (FOV) could be reconstructed (see Fig. S1d). The data from the same FOV, as in Fig. S1d, was used for the analyses presented in the main text and the SI.

**Temporal evolution of a cell monolayer:** First, we explore some general features of the temporal evolution of the MDCK monolayer using the experimentally generated trajectories[1]. Although the MDCK monolayer is confluent initially, it does not stay in a stationary state but evolves with time, exhibiting rich dynamics driven by cell division and apoptosis.

*Cell divisions and apoptosis:* The number of cell divisions (see Fig. S1c) between two frames (one minute interval) is quantified in Fig. S2a (see the green circles). Cell apoptosis/extrusion events are also shown in the same figure by yellow stars. These two numbers fluctuate between 0 and 12. Therefore, the confluent cell monolayer is dynamic that undergoes continuous cell division and apoptosis, which generate active forces leading to the collective cell motions[4–6]. The mean value for the number of new cells (5.14) is a slightly higher than the dead cells (5.05). Hence, the number of cells in the FOV increases with time, as shown in Fig. S2b. A power-law relation describes the cell number as a function of time that is shown by the green solid line and the expression is given in the figure caption.

*Cell size distributions:* Due to cell growth, division and apoptosis, the cell size is not uniformly distributed. In order to calculate the distribution of cell sizes, we used Voronoi tessellation of the positions of all the cell nuclei in the FOV (Fig. S1d). The Voronoi cells at two different time frames are shown in Figs. S3a-b, and the area of each cell is color coded as indicated in the bar on the right side of the figures. There is a large variation (from 100 to 400 $\mu m^2$) in the cell area. The cell area distributions, $P(A)$, at early (sampled over 20 images from $t$ = 180 to 199 minutes) and later (1350-1369 minutes) times are illustrated in Fig. S3c (without considering the boundary Voronoi cells). A Gaussian distribution (see the solid lines and the functional forms shown in Fig. S3c) describes the $P(A)$ reasonably well.

The mean value of the cell area decreases from 231 $\mu m^2$ at early times to 187 $\mu m^2$ at later times during a 24-hour period in the experiments. This is an indication of cell jamming[7,8].

*Cell density and velocity:* As the number of cells increases (see Fig. S2b) in the same FOV, so does the cell density (see Fig. S4a, defined as the ratio of the cell number and the area in the FOV). A similar power-law relation is found for the cell density and the cell number (see the solid lines and functional forms shown in Fig. S2b and Fig. S4a). The mean velocity, defined in the main text above Eq. (1) with $\delta t = 10$ min) of cells, decreases with time (see the green line in Fig. S4b). Therefore, the overall motility of cells is inhibited as the cell density increases, even though the frequencies of active cell division and apoptosis are nearly constant (see Fig. S2a).

*Distribution of cell coordination number:* The number of neighbors (coordination number, $Z$) per cell in MDCK monolayer varies from cell to cell (see Figs. S3a-b) because of the generation of active forces. From the Voronoi tessellation (Fig. S3), we calculated the cell coordination number distribution, $P(Z)$, for the MDCK cell monolayer (see the histograms in blue in Fig. S5). Each histogram is calculated using ten time frames from the experiments[1]. The value of $Z$ varies from 4 to 9, and its distribution is well described by a Gaussian distribution with a mean value around 6 (see the solid lines), which remains robust during the 24 hours experiment (see Figs. S5a-b). The distribution, $P(Z)$, obtained from our simulations agrees well with the experimental results (see the histograms and also the solid lines in Fig. S5). Therefore, a predominantly hexagonal pattern can be driven purely by cell division and apoptosis, which leads to an emergent phenomena of geometric order, similar to that found in proliferating metazoan epithelia[9].

**Displacement distribution of cells:** We calculated the displacement distributions ($P(\Delta x)$, $P(\Delta y)$) of cells using the trajectories from the MDCK cell experiments and our simulations (see Fig. S6). Dramatic deviations from the Gaussian distribution (see the green solid lines) are found in both experiments and simulations.

**T2 transition induced by cell apoptosis:** The T1 transition induced by cell divisions is illustrated in Figs. 2e-g in the main text. We also found evidence for another type of topological change, the T2 transition, caused by cell apoptosis/extrusion from the confluent monolayer (see Fig. S8). Through Voronoi tessellation, using the cell nuclei position data, we found that cell 0 (the one shown in red) is extruded from the cell monolayer at a later time (see Fig. S8b), leading to the formation of new neighborhood constituting cells $1-6$. For completeness, we should point out that T1 and T2 transitions are well known in the dynamics of foams[11] where connections to cells was also explained.

**Velocity autocorrelation at different $\delta t$:** We showed that the velocity autocorrelation function exhibits long time tail (LTT) behavior in the main text (see Fig. 4c). To ascertain whether LTT depends on the time interval, $\delta t$, we calculated velocity autocorrelation of MDCK cells[1] at several values of $\delta t$ (2, 4, and 10 minutes). The result is robust (see the different symbols and the dash-dotted lines in Fig. S10).

**Energy spectrum:** The kinetic energy $E$ is given by,

$$E = \frac{1}{2}\int |\mathbf{v}(\mathbf{r})|^2 d\mathbf{r} = \int E(\mathbf{k})d\mathbf{k}, \tag{1}$$

where $\mathbf{v}(\mathbf{r})$ is the cell velocity at position $\mathbf{r}$, and $k = |\mathbf{k}|$ is the wave number. To calculate the energy spectrum $E(k)$ at a time point, we first use the digital particle image velocimetry[10] to get the velocity field $\mathbf{v}(\mathbf{r})$ (using two images with $\delta t = 10$ minutes) at regular grid points. Then, we used the fast Fourier transform (FFT) to convert the velocity field $\mathbf{v}(\mathbf{r})$ to $\tilde{\mathbf{v}}(\mathbf{k})$. Finally, the energy spectrum $E(k)$ is calculated using,

$$E(k) = \frac{kA_0}{4\pi}\sum \tilde{\mathbf{v}}^*(\mathbf{k}) \cdot \tilde{\mathbf{v}}(\mathbf{k}), \tag{2}$$

where $\tilde{\mathbf{v}}^*(\mathbf{k})$ is the complex conjugate of $\tilde{\mathbf{v}}(\mathbf{k})$ and $A_0$ is the area of the FOV. Two examples for $E(k)$ are illustrated here: one is the Fig. 4e in the main text and the other is shown in Fig. S11. In each figure, we calculated the $E(k)$ at six different time frames. The dependence of $E(k)$ on $k$ shows that there are two scaling regimes (see the solid lines and also the values of the slope). The two solid lines give the best power-law fit over the data (all six time frames) for different regimes of the wavevector $k$.

**Agent-based simulations:** We use a two dimensional (2D) version of the three dimensional (3D) agent-based model[12,13]. In 2D, the cells are discs. We just give a sketch of the model because additional details can be found elsewhere[12]. We use a deformable disc to represent each cell. The cell area can increase at a constant rate ($r_A$) as long as the pressure, $p$, it experiences due to contact with the neighboring cells is less than a preassigned critical value $p_c$[12]. If $p$ exceeds $p_c$, the cell enters a dormant state, stops growing unless the normal pressure is below $p_c$, which could occur as the system evolves. The use of $p_c$ accounts for cell proliferation inhibition by a mechanical feedback mechanism[14–18]. After reaching the fixed mitotic radius $R_m$, with $p < p_c$, a cell divides into two with identical radius, $R_d = R_m 2^{-1/2}$ ($\pi R_m^2 = 2\pi R_d^2$). The new cells are placed randomly in space (the mother cell occupied before division) at a center-to-center distance $d = 2R_m(1 - 2^{-1/2})$. The cell area grows at a rate $r_A = \pi R_m^2/(2\tau_{min})$ ($\tau_{min}$ is the cell cycle time). We update the cell radius ($R$) from a Gaussian distribution with the mean value $\dot{R} = r_A/(2\pi R)$.

The elastic force $F_{ij}^{el}$ between two cells of radii $R_i$ and $R_j$ are modeled by Hertzian contact mechanics[19–21],

$$F_{ij}^{el} = \frac{h_{ij}^{3/2}(t)}{\frac{3}{4}\left(\frac{1-\nu_i^2}{E_i} + \frac{1-\nu_j^2}{E_j}\right)\sqrt{\frac{1}{R_i(t)} + \frac{1}{R_j(t)}}}, \tag{3}$$

where $E_i$ and $\nu_i$ are the elastic modulus and Poisson ratio of cell $i$, respectively. The overlap $h_{ij}$ between the two cells is defined as $\max[0, R_i + R_j - |\vec{r}_i - \vec{r}_j|]$ with $|\vec{r}_i - \vec{r}_j| \equiv r_{ij}$, being the center-to-center distance of the cells. In the simulations, we use $E_i$ and $\nu_i$ to be independent of $i$. Cell-cell adhesive interaction, $F_{ij}^{ad}$, is mediated by receptor and ligand proteins on the cell membrane which is calculated using[19],

$$F_{ij}^{ad} = L_{ij} f^{ad} \frac{1}{2}(c_i^{rec} c_j^{lig} + c_j^{rec} c_i^{lig}), \tag{4}$$

where the $c_i^{rec}$ ($c_i^{lig}$) is the receptor (ligand) concentration, which is considered to be constant here, and $f^{ad}$ is a coupling constant used to rescale the adhesion force. The length of the contact line, $L_{ij}$ between two cells, is given by $L_{ij} = \sqrt{|4r_{ij}^2 R_i^2 - (r_{ij}^2 - R_j^2 + R_i^2)^2|}/r_{ij}$. The total force on the $i^{th}$ cell is the sum over its nearest neighbors ($NN(i)$),

$$\vec{F}_i = \Sigma_{j\epsilon NN(i)}(F_{ij}^{el} - F_{ij}^{ad})\vec{n}_{ij}. \tag{5}$$

where $\vec{n}_{ij}$ is the unit vector pointing from the center of cell $j$ to the center of cell $i$.

The spatial dynamics of the cell is calculated by integrating the equation of motion for a cell of mass $m_i$,

$$m_i \ddot{\vec{r}}_i = \vec{F}_i(t) - \gamma_i \dot{\vec{r}}_i(t), \tag{6}$$

where $\gamma_i$ is the modified mobility coefficient of cells with $\gamma_i = \gamma_i^{visc} + \gamma_i^{ad}$. The first term comes from the contribution of cell to extracellular matrix (ECM) friction ($\eta r_i$, with $\eta$ the mobility constant) and the second term describes the cell-to-cell friction with,

$$\begin{aligned} \gamma_i^{ad} = \quad & \gamma^{max} \Sigma_{j\epsilon NN(i)} [L_{ij} \frac{1}{2}(1 + \frac{\vec{F}_i \cdot \vec{n}_{ij}}{|\vec{F}_i|}) \times \\ & \frac{1}{2}(c_i^{rec} c_j^{lig} + c_j^{rec} c_i^{lig})] \,. \end{aligned} \tag{7}$$

Because Reynolds number for cells in a tissue is low[22], the inertial term in Eq. (6) can be neglected[19]. Therefore, the equation of motion becomes,

$$\dot{\vec{r}}_i = \frac{\vec{F}_i}{\gamma_i}. \tag{8}$$

We calculated the pressure $p$ as in previous studies (Eq. (8) in Ref.[12]) except the contact line length, $L_{ij}$, is used instead of the contact area, $A_{ij}$, between two cells. We employed periodic boundary conditions instead of the free boundary for a growing tumor spheroid[12,13].

We start from 300 cells that are randomly distributed in a box with the size $340\mu m \times 340\mu m$, similar to the one in the FOV in the experiments[1] (see Fig. 1 in the main text). The parameters in the agent-based model used in the simulations are given in Table I. Note that the ratio of the rate of apoptosis to cell division (0.54) is comparable to our estimation using the experimental data.

TABLE I: The parameters used in the simulation.

| **Parameters** | **Values** | **References** |
|---|---|---|
| Time step ($\Delta t$) | 10 s | [12] |
| Critical Radius for Division ($R_m$) | 11 $\mu$m | This paper |
| Mobility constant ($\eta$) | 0.1 kg/($\mu$m s) | This paper |
| Benchmark Cell Cycle Time ($\tau_{min}$) | 54000 s | [24–26] |
| Adhesive Coefficient ($f^{ad}$) | $10^{-4}\mu$N/$\mu$m | [19] |
| Mean Cell Elastic Modulus ($E_i$) | $10^{-3}$MPa | [23] |
| Mean Cell Poisson Ratio ($\nu_i$) | 0.5 | [19] |
| Death Rate ($b$) | $10^{-5}$s$^{-1}$ | This paper |
| Mean Receptor Concentration ($c^{rec}$) | 1.0 (Normalized) | [19] |
| Mean Ligand Concentration ($c^{lig}$) | 1.0 (Normalized) | [19] |
| Adhesive Friction $\gamma^{max}$ | $10^{-4}$kg/($\mu$m s) | [12] |
| Threshold Pressue ($p_c$) | $5 \times 10^{-3}\mu$N/$\mu$m | This paper |

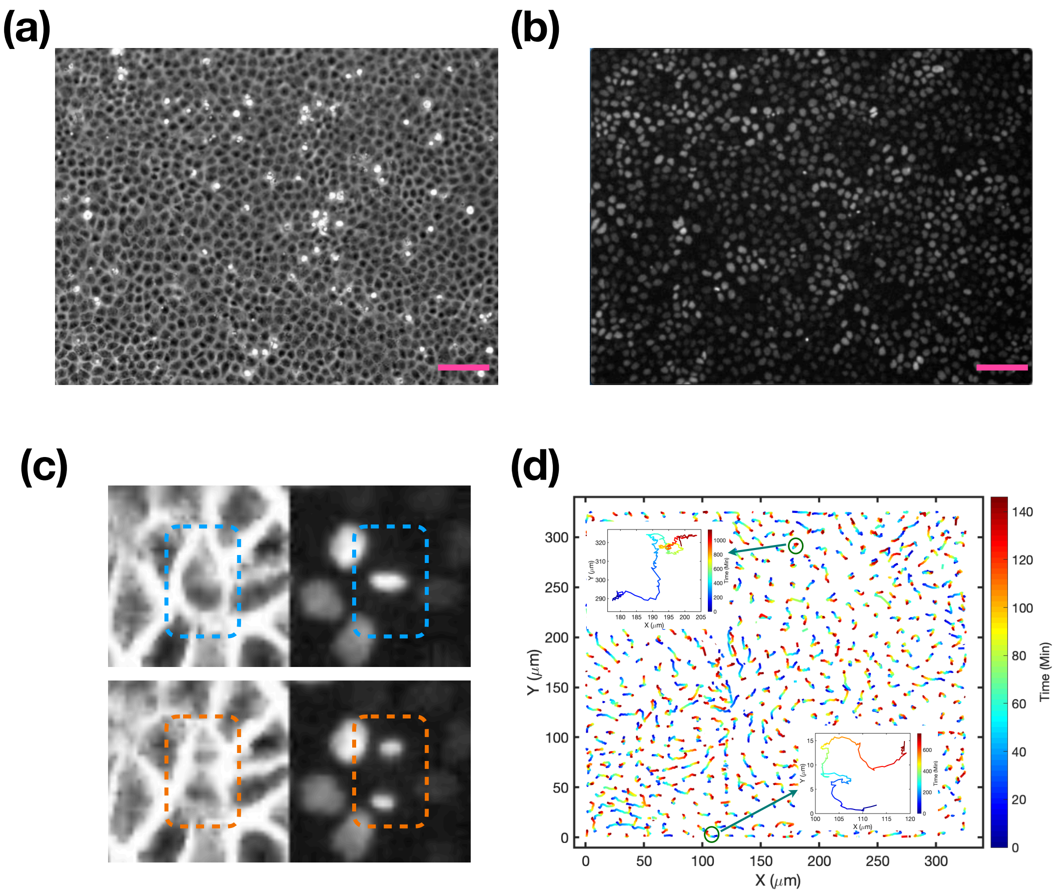


FIG. S1: **MDCK monolayer.** (a) A representative image of a confluent MDCK monolayer (t = 0 min) using the phase-contrast microscopy (taken from experiments by Fabio et al.[1]), which highlights the cell membrane. (b) Same as (a) except the image is taken using the wide-field fluorescence microscopy[1], where the cell nuclei are highlighted. The size of the images in (a)-(b) is (867 × 660) $\mu m^2$, and the pink bars correspond to 100 $\mu m$. (c) A cell division event[1]. Upper (lower) panel: before (after) a cell division. The left panel are images from the phase-contrast microscopy. Right panel: images from the wide-field fluorescence microscopy for EGFP-H2B expressing cells. The size of the image is (60 × 60) $\mu m^2$. (d) The cell trajectories in one field of view (FOV) are shown over the first 150 minutes. The trajectory is color coded by time (see the color bar on the right of the figure). The insets display two representative cell trajectories over 24 hours. Heterogeneity in the dynamics and swirl patterns are evident.

(a)

(b)

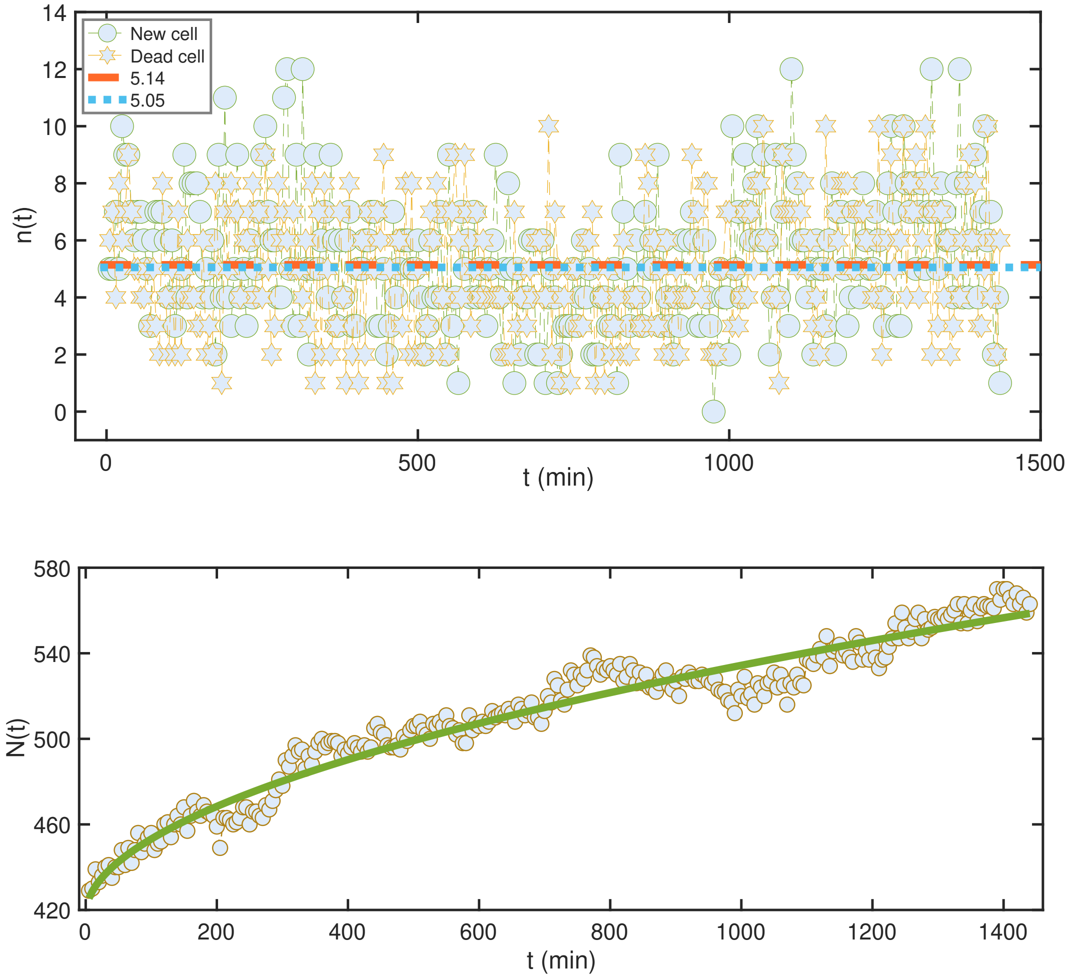


FIG. S2: **Cell number dynamics.** (a) The number of new and dead or extruded cells ($n(t)$) in one FOV (see Fig. S1d) over the duration of the experiment. The red dashed and blue dotted lines are the mean values for the new and dead/extruded cells, respectively. (b) The temporal evolution of the cell number ($N(t)$ in the same FOV as (a). The green curve is a power law ($N(t) = 418 + 3.2t^{0.5}$) fit of the experimental data in circles.

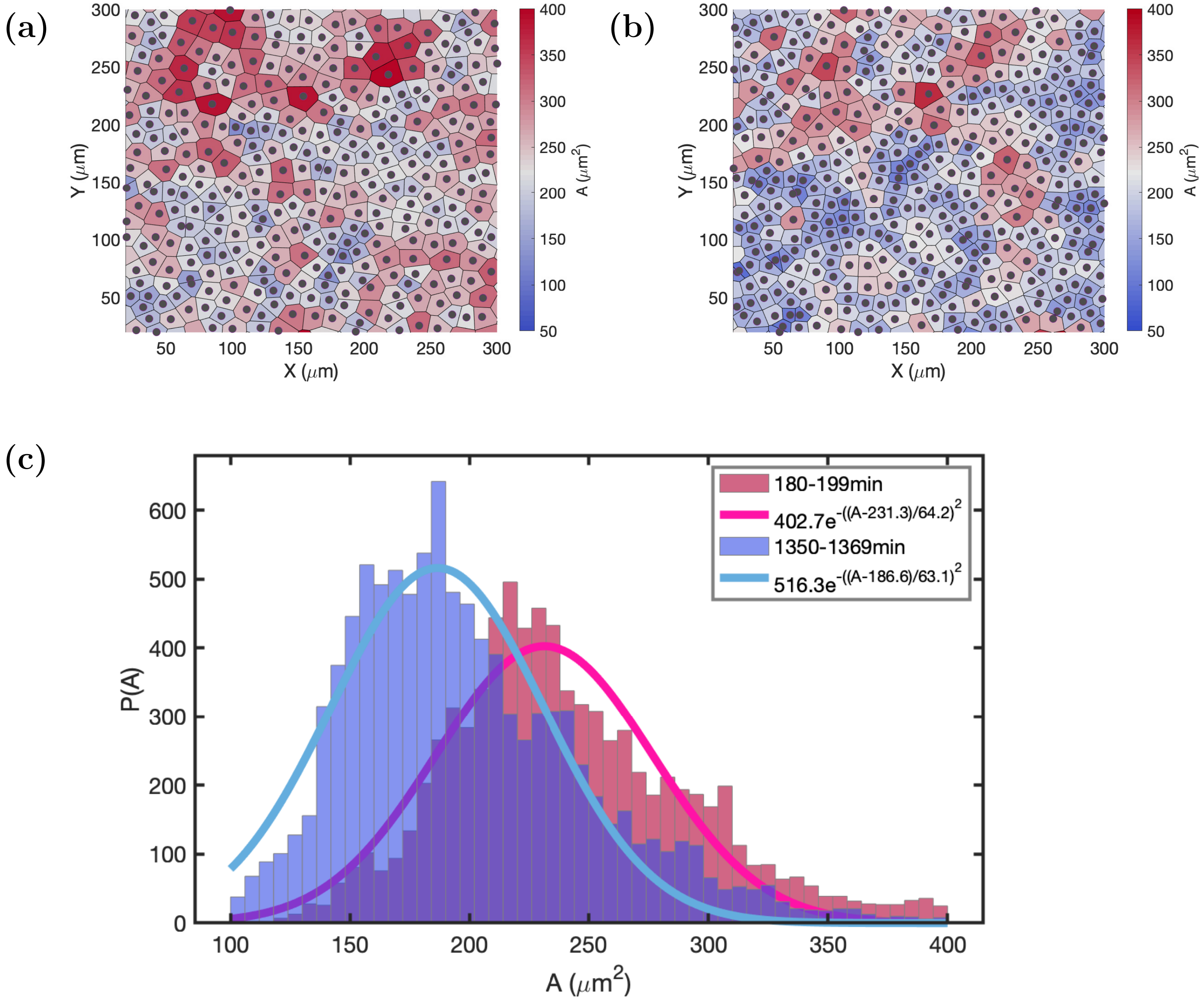


FIG. S3: **Cell size distributions of epithelial monolayers.** (a) A representative image of a confluent cell monolayer at an early time ($t$ = 180 minute). (b) Same as (a) except $t$ = 1350 minutes in the experiments. In (a) and (b), the size of each Voronoi cell is color coded according to the area of each cell (see the bar on the right of the figure). (c) The distribution $P(A)$ of the cell area ($A$) between $t$ = 180 to $t$ = 199 minutes (blue), and between $t$ = 1350 and $t$ = 1369 minutes (pink). The solid lines show the best fit to the data. The functional forms are in the inset.

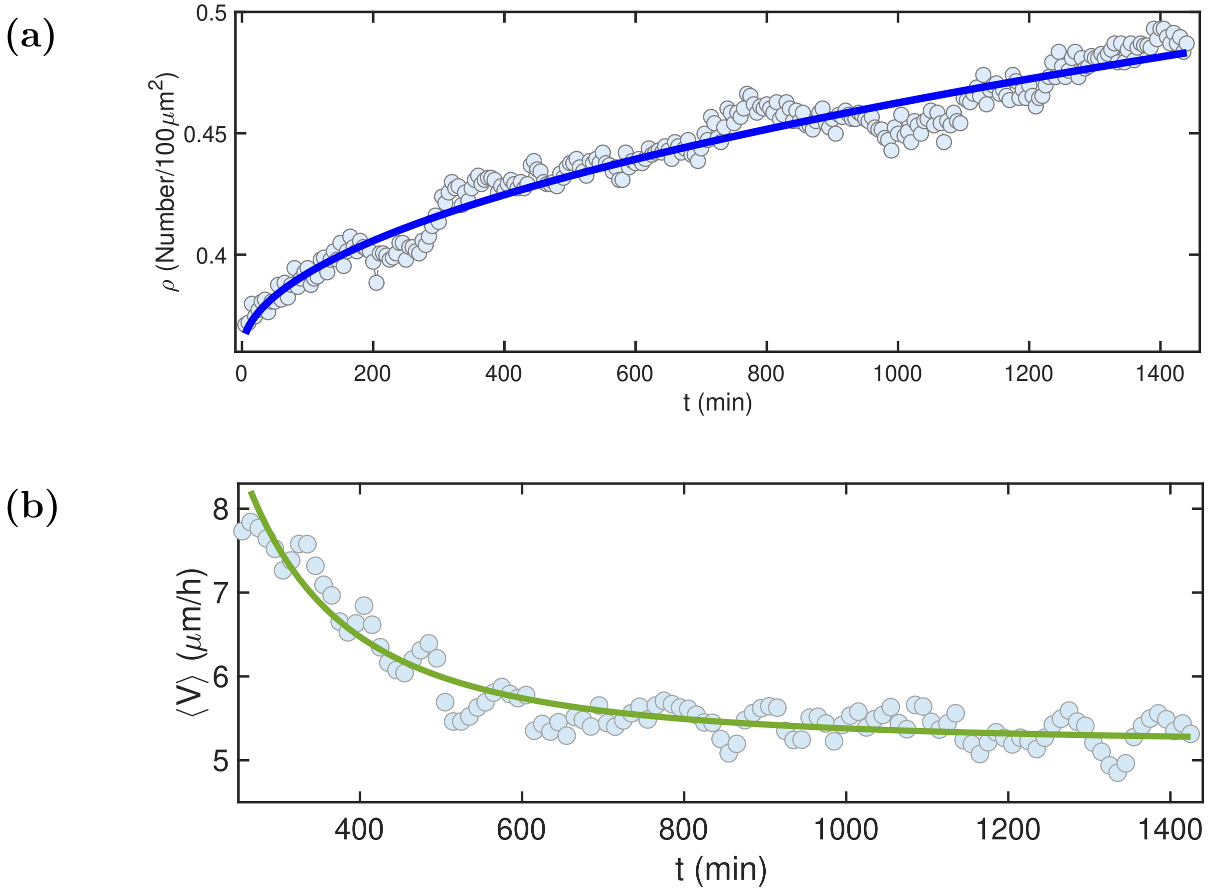


FIG. S4: **Cell number density ($\rho$) and mean velocity ($\langle V \rangle$).** (a) The evolution of $\rho$ in unit of number per 100 $\mu m^2$. The blue line is the power law fit ($\rho = 0.36 + 0.003t^{0.5}$) of the experimental data, shown in circles. (b) The mean velocity of cells as a function of time. Each circle represents the mean velocity of all the cells in ten time frames. The green line, which is the power law fit ($\langle V \rangle = 5.2 + 3.3 \times 10^5 t^{-2.05}$) of the experimental data, shows a decrease as the cell density increases, which is an indication of cell jamming.

(a)

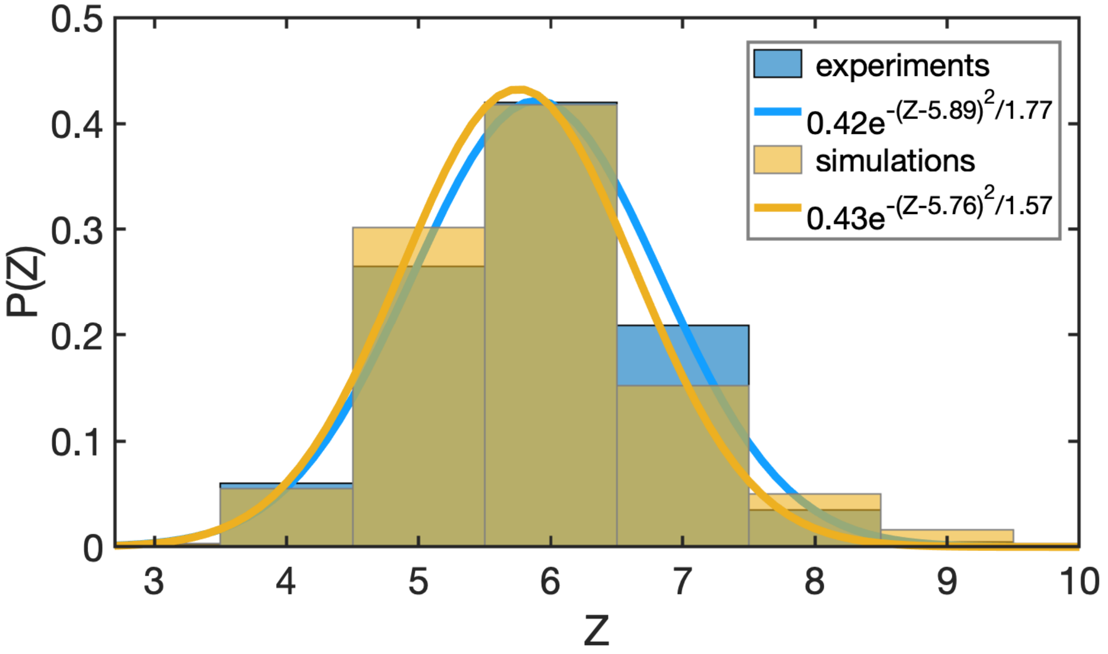


(b)

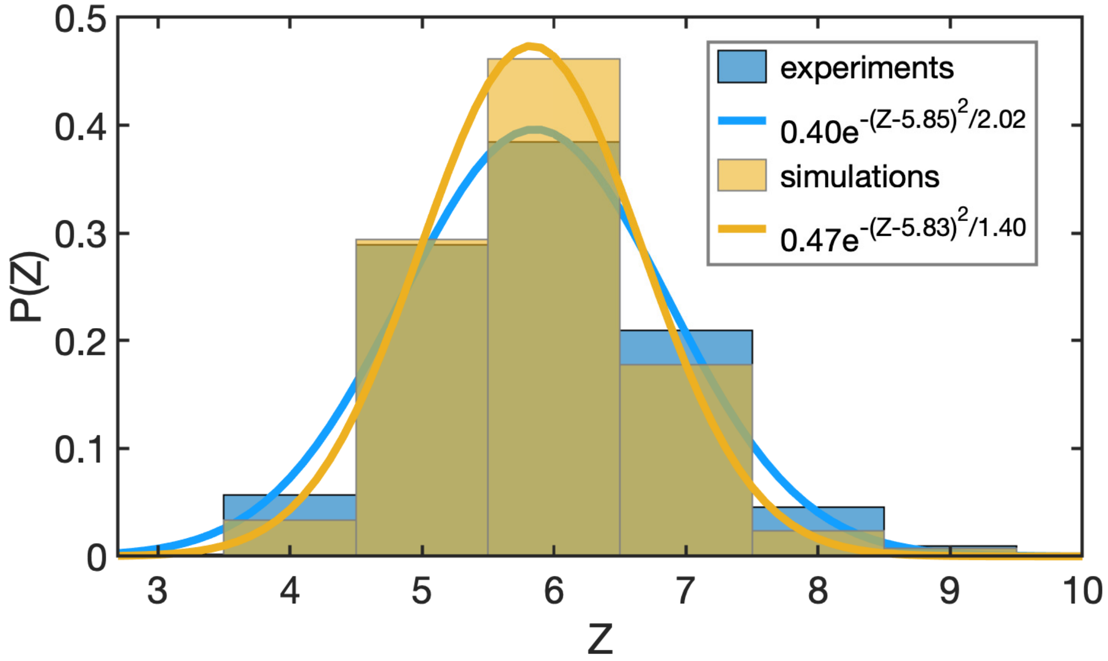


FIG. S5: **Distribution of cell coordination number ($Z$).** (a) The distribution ($P(Z)$) of $Z$ between $120-129$ minutes. (b) Same as (a) except the time interval $1000-1009$ minutes. The blue (brown) histograms are calculated from experiments (simulations). The solid blue and brown lines are Gaussian fits to the experimental and simulation results, respectively.

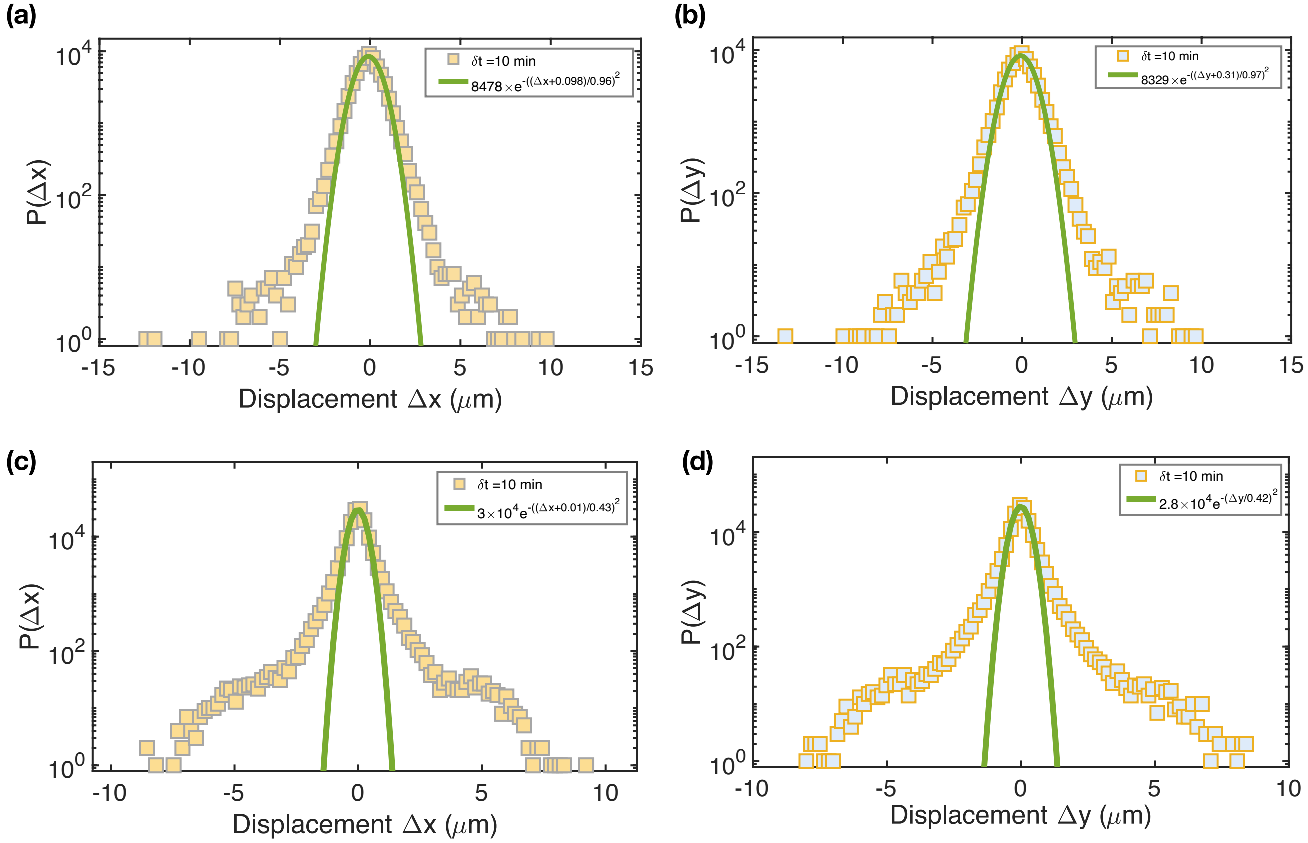


FIG. S6: **Anomalous diffusion.** (a-b) Distribution (P(Δx), P(Δy)) of cell displacement at the time interval, $\delta t$, of 10 minutes calculated for all the cells over the whole experiment. The green lines show Gaussian distributions. (c-d) Same as (a-b), except the distributions are obtained from simulations.

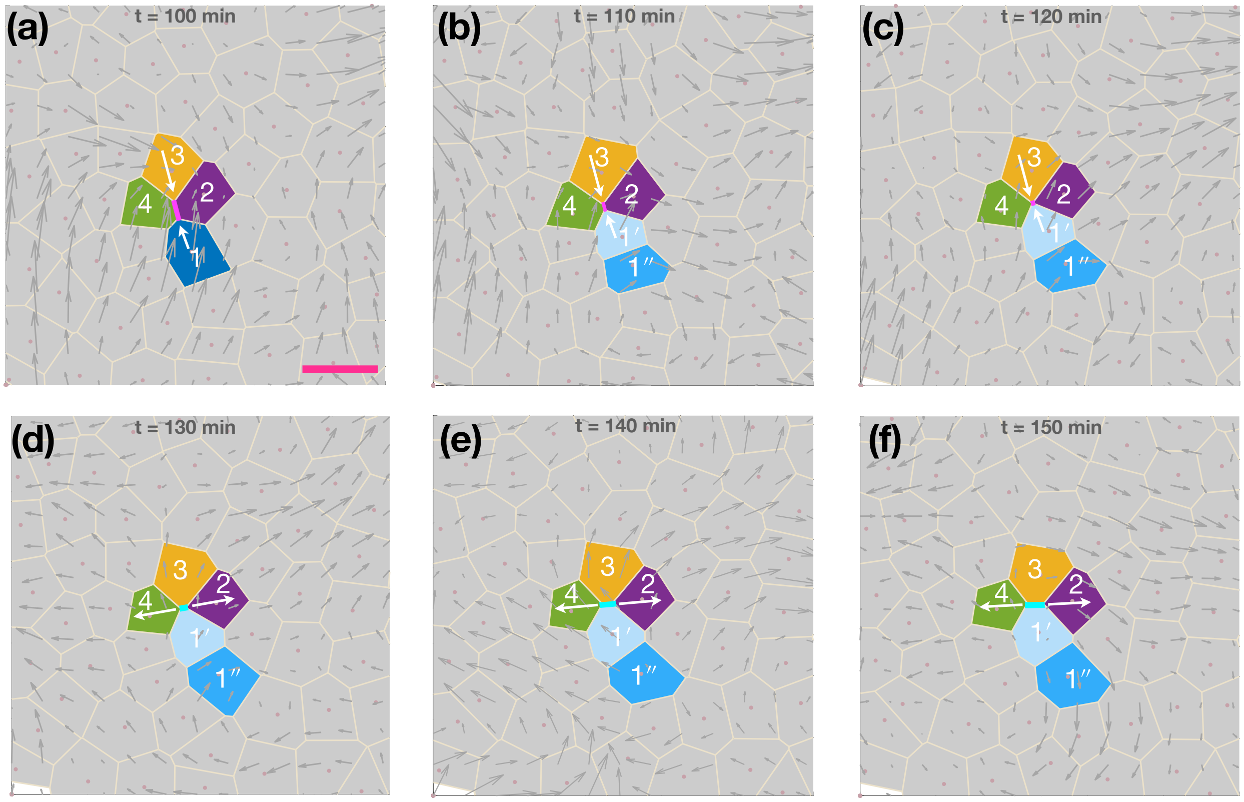


FIG. S7: **T1 topological changes.** Same as Figs. 2e-g in the main text, except more time frames are included. There is a T1 transition, accompanied by one vortex formation, which is induced by a single cell division (cell 1 divided into cells $1'$ and $1''$) . The pink bar in (a) corresponds to 20 $\mu m$. The velocity vector field is illustrated by grey arrows.

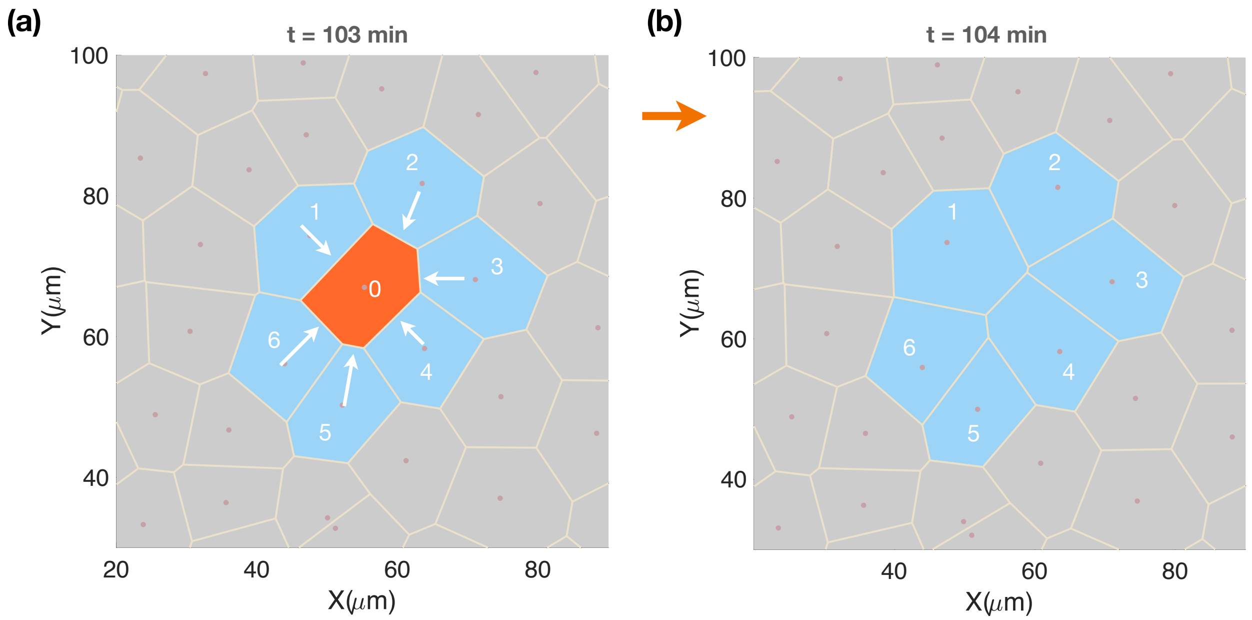


FIG. S8: **Cell apoptosis (extrusion) induces T2 transition.** Cell 0 (red), surrounded by cells $1-6$ (blue) at time $t = 103$ min (a) is removed from the cell monolayer at time $t = 104$ min due to apoptosis/extrusion process, which results in the topological T2 transition (shown in (b)).

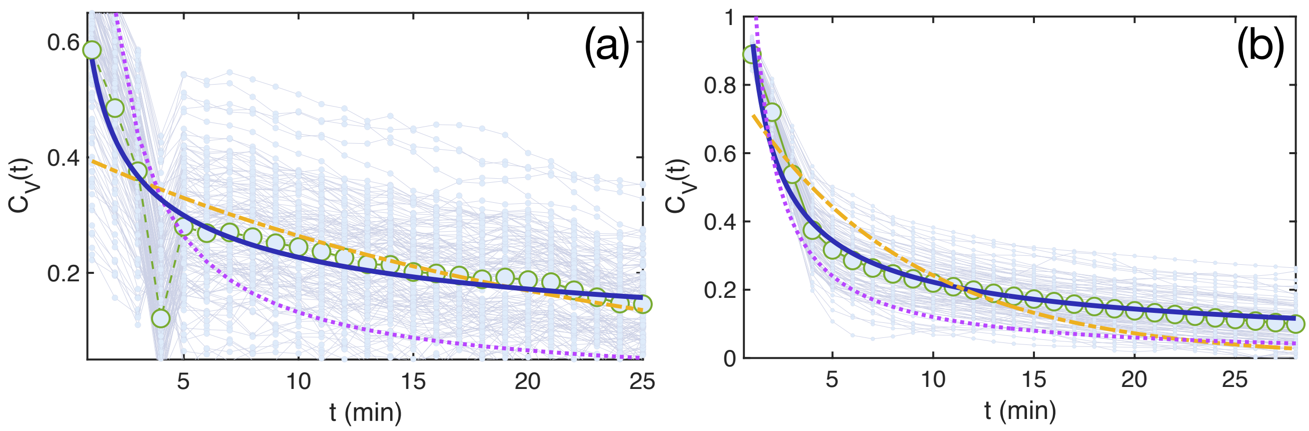


FIG. S9: **Velocity autocorrelation:** Same as Figs. 4(c-d) in the main text. (a) Experimental, and (b) simulation data fit using different functions. The blue solid line shows a power-law fit ($t^{-\beta}$ with $\beta \approx 0.4, 0.6$ as listed in Figs. 4(c-d)). The yellow dash-dotted line indicates an exponential fit, and the violet dotted line shows a fit using $t^{-1}$, the expected behavior for 2D liquids. It is clear that the best fit is obtained using $t^{-\beta}$ (with $\beta < 1$).

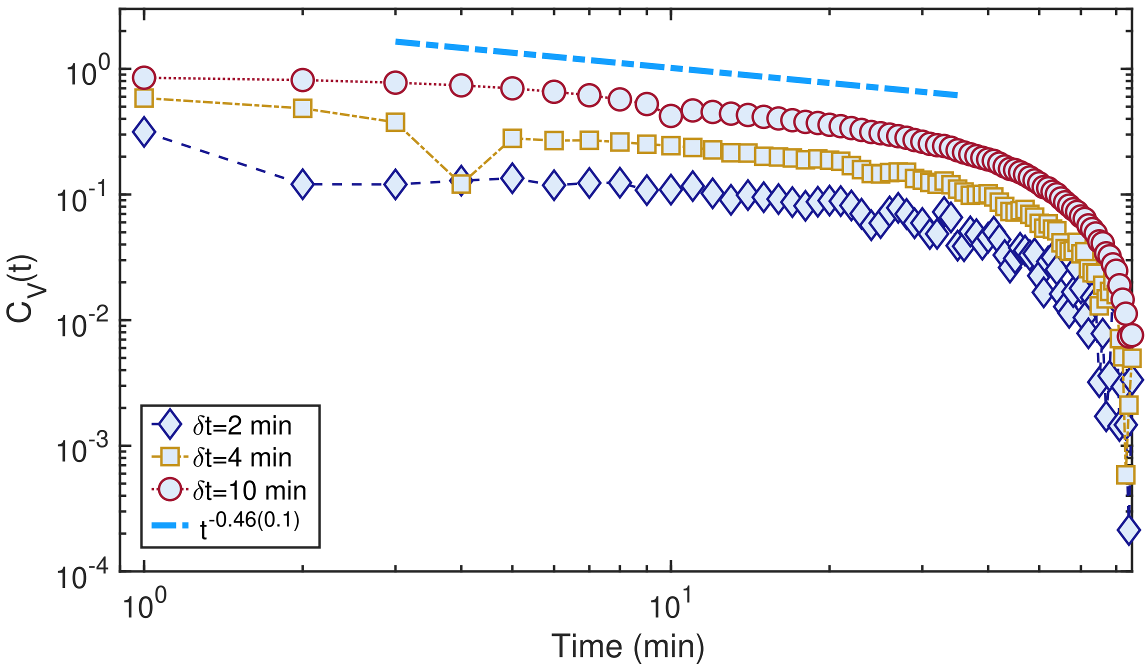


FIG. S10: **Velocity autocorrelation at different** $\delta t$**:** (a) Same as Fig. 4(c) in the main text. We calculated $C_v(t)$ with $\delta t = 2$, 4, and 10 minutes. The slope of the dashed-dotted line is shown in the box with errors listed in the bracket. Thus, variations in $\delta t$ do not change the emergence of long time tails.

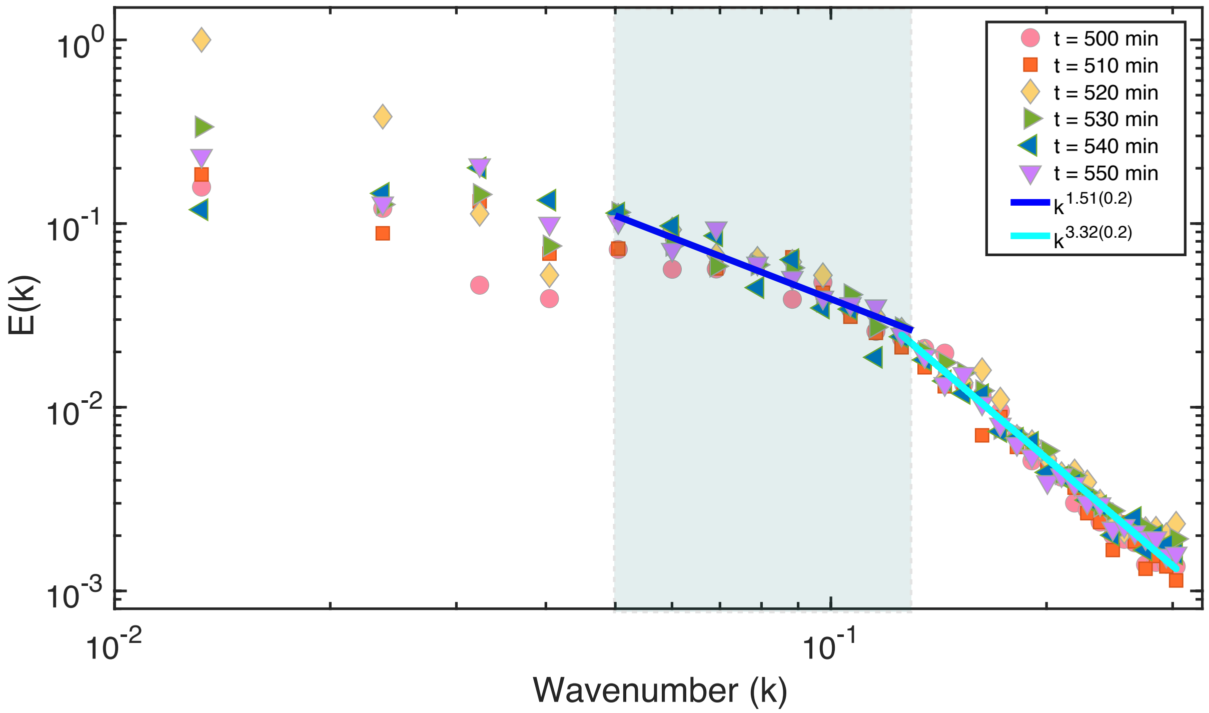


FIG. S11: **The energy spectra E(k) at different times.** Same as Figure 4(e) in the main text, except at different times ($t = 500 - 550$ minutes). The values of the slope of the solid lines are shown in the upper right corner with errors listed in the bracket. The unit for the wave number ($k$) is $\mu\mathrm{m}^{-1}$, and $E(k)$ ($\mu\mathrm{m}^3/\mathrm{s}^2$) is scaled by the maximum value.
